# Microglia-dependent LPS preconditioning prevents neuroinflammation-induced behavioral deficits in male mice

**DOI:** 10.1101/2025.10.10.681580

**Authors:** Minori Koga, Hiroyuki Nakashima, Masafumi Saito, Mayumi Sato, Ryuichi Nakagawa, Takanobu Sato, Fumiho Asai, Tamae Ishii, Manabu Kinoshita, Masanori Nagamine, Hiroyuki Toda

## Abstract

Neuroinflammation contributes to psychiatric disorders, but preventive strategies targeting brain immune cells remain unexplored. Here we demonstrate that low-dose lipopolysaccharide (LPS) preconditioning prevents systemic inflammation-induced behavioral abnormalities through microglia-dependent mechanisms in male mice. Mice received preconditioning with 0.2 mg/kg LPS or saline for two consecutive days, followed by high-dose LPS challenge (5 mg/kg) or saline seven days later. Behavioral assessment revealed that preconditioning specifically prevented social preference deficits induced by systemic inflammation (preference score: -0.49±0.19 vs 0.14±0.10, p<0.01), while showing limited effects on locomotor activity and depression-like behaviors. Additionally, LPS preconditioning prevented anxiety-like behavior in a chronic corticosterone model and attenuated hippocampal inflammatory gene expression. Immunohistochemical analysis demonstrated that preconditioning suppressed microglial activation in hippocampal CA1 region, particularly reducing PBR/IBA1 ratio (37.5±2.4% vs 27.6±2.8%, p<0.01), with less pronounced effects in CA3. Critically, pharmacological microglial depletion using PLX3397 during the preconditioning period completely abolished these protective effects, establishing the causal role of microglia. Flow cytometric analysis revealed preconditioning-induced shifts in brain macrophage subpopulations defined by TMEM119 and CD45 expression patterns. Transcriptomic profiling identified subpopulation-specific responses, with one subset showing LPS-response pathway enrichment despite minimal gene expression changes, while another displayed extensive but functionally non-specific transcriptional alterations. These findings establish microglial preconditioning as a novel preventive strategy for neuroinflammation-induced social behavioral deficits and suggest potential therapeutic applications for psychiatric disorders involving neuroinflammatory components.

## Introduction

Psychiatric disorders represent a major global health burden, and elucidating their pathogenic mechanisms while developing preventive and therapeutic strategies remains an urgent challenge. Recently, the "neuroinflammation hypothesis" has gained attention, proposing that neuroinflammation contributes to the pathogenesis of psychiatric disorders including depression and schizophrenia ^1-3^. Evidence from patient brains, animal models, and epidemiological studies has demonstrated elevated inflammatory cytokines, activation of brain glial cells, and correlations between blood inflammatory markers and symptoms ^4-6^, suggesting that abnormalities in brain immunity may be risk factors for psychiatric symptom development.

Observational and PET imaging studies have reported increased TSPO expression and neuroinflammatory markers in psychiatric patients ^7, 8^. Additionally, sepsis survivors often exhibit subsequent depressive and anxiety-like psychiatric symptoms ^9, 10^. Furthermore, associations between inflammatory gene polymorphisms and depression risk ^11^, as well as correlations between peripheral inflammatory markers and symptom severity or treatment resistance, have been demonstrated ^12, 13^. These findings support the possibility that inflammation is not merely an epiphenomenon but contributes to pathogenesis itself. Nevertheless, most previous studies have only demonstrated correlational relationships between inflammation and psychiatric symptoms, with few clearly establishing causal relationships.

As interventions targeting neuroinflammation, studies have aimed to prevent or alleviate psychiatric symptoms through adjunctive anti-inflammatory drugs. While COX-2 inhibitors such as celecoxib have been reported to reduce depressive symptoms or prevent onset ^14, 15^, other reports show no effect ^16, 17^, demonstrating inconsistent efficacy ^16, 17^. This inconsistency suggests that the pathophysiology of psychiatric disorders is diverse and not solely dependent on conventional inflammatory cascades. Moreover, previous studies based on the neuroinflammation hypothesis have predominantly focused on inflammatory cytokines and mediators, with limited investigation into the significance and roles of brain immune cells themselves.

Microglia are the primary immune cells of the central nervous system, producing inflammatory cytokines while also contributing to neural homeostasis and behavioral control ^18^. Although microglial involvement in psychiatric symptom development is suspected, empirical research on how they actually contribute to symptom formation or prevention remains scarce ^19^. Particularly, whether psychiatric symptoms can be prevented by pre-emptively regulating the immune system, and evaluation of the significance and efficacy of such preventive interventions, remains largely unexplored.

Recent technological advances have revealed that microglia are not a single homogeneous cell population but comprise subpopulations with different phenotypes and functions ^20^. We previously reported that brain CD11b-positive cells can be classified into multiple subpopulations based on TMEM119 and CD45 expression patterns in a chronic corticosterone-induced HPA axis hyperactivity model (Nakagawa et al., 2025). This study identified increased TMEM119-high CD45-intermediate microglia correlating with anxiety-like behavior, suggesting involvement of specific microglial subtypes in stress-related behavioral abnormalities. However, the roles these subpopulations play in neuroinflammation control and preventive interventions remain unclear.

The hippocampus plays a crucial role as a neural substrate for social behavior ^21^. Specifically, the CA3 region is important for pattern separation of novel stimuli and social memory formation ^22^, while the CA1 region is known to be involved in information integration and memory retrieval ^23^. In neuroinflammation-induced social behavioral impairments, the degree and pattern of microglial activation in these regions may differ, though details remain unclear.

This study focused on lipopolysaccharide (LPS) preconditioning/tolerance, which is induced by repeated injections with low-dose LPS from the perspective of preventively controlling microglial activity itself. LPS, a well-characterized pathogen-associated molecular pattern (PAMP), is known to directly activate microglia via Toll-like receptor 4 (TLR4) ^24, 25^. Given this property, we hypothesized that prior exposure to a low dose of LPS may modulate the subsequent activation profile of microglia. Preconditioning represents a state of immune tolerance, in which prior mild stimulation attenuates excessive inflammatory responses. While this concept has been extensively applied in models of cerebral ischemia ^26^, its potential to prevent inflammation-induced behavioral abnormalities has not been fully explored.

We evaluated whether low-dose LPS preconditioning could suppress behavioral abnormalities caused by LPS-induced systemic inflammation^27^ and verified the necessity of microglia for this effect. Furthermore, using brain macrophage subpopulations we previously identified ^28^, we analyzed in detail the quantitative changes and transcriptional profile alterations in each subpopulation induced by preconditioning. Through these investigations, we aimed to elucidate microglial subpopulation-specific immune regulatory mechanisms and obtain findings that could serve as the foundation for novel preventive and therapeutic strategies for neuroinflammation-related psychiatric disorders.

## Material and Methods

### Animal

Male Institute of Cancer Research (ICR; CD-1) mice, aged 10 weeks, were purchased from Japan SLC (Hamamatsu, Japan) and maintained in a temperature (22–24 °C) and humidity (50%) conditioned room with a 12:12 hour light-dark cycle, with lights on at 7:00 AM. Food (CE-7; Clea Japan, Tokyo, Japan) and water were supplied *ad libitum*. Mice were housed in standard cages measuring 225×338×140 mm with sawdust. A maximum of five mice were housed per cage. All experiments were performed in strict accordance with ARRIVE guidelines and approved by the local Animal Investigation Committee of the National Defense Medical College (Approval number: 22015, 22042, and 23084)

### Experimental design and pharmacological administration

Four independent experiments were conducted in this study. The experimental timelines are illustrated in Figure 1.

**Figure 1.**
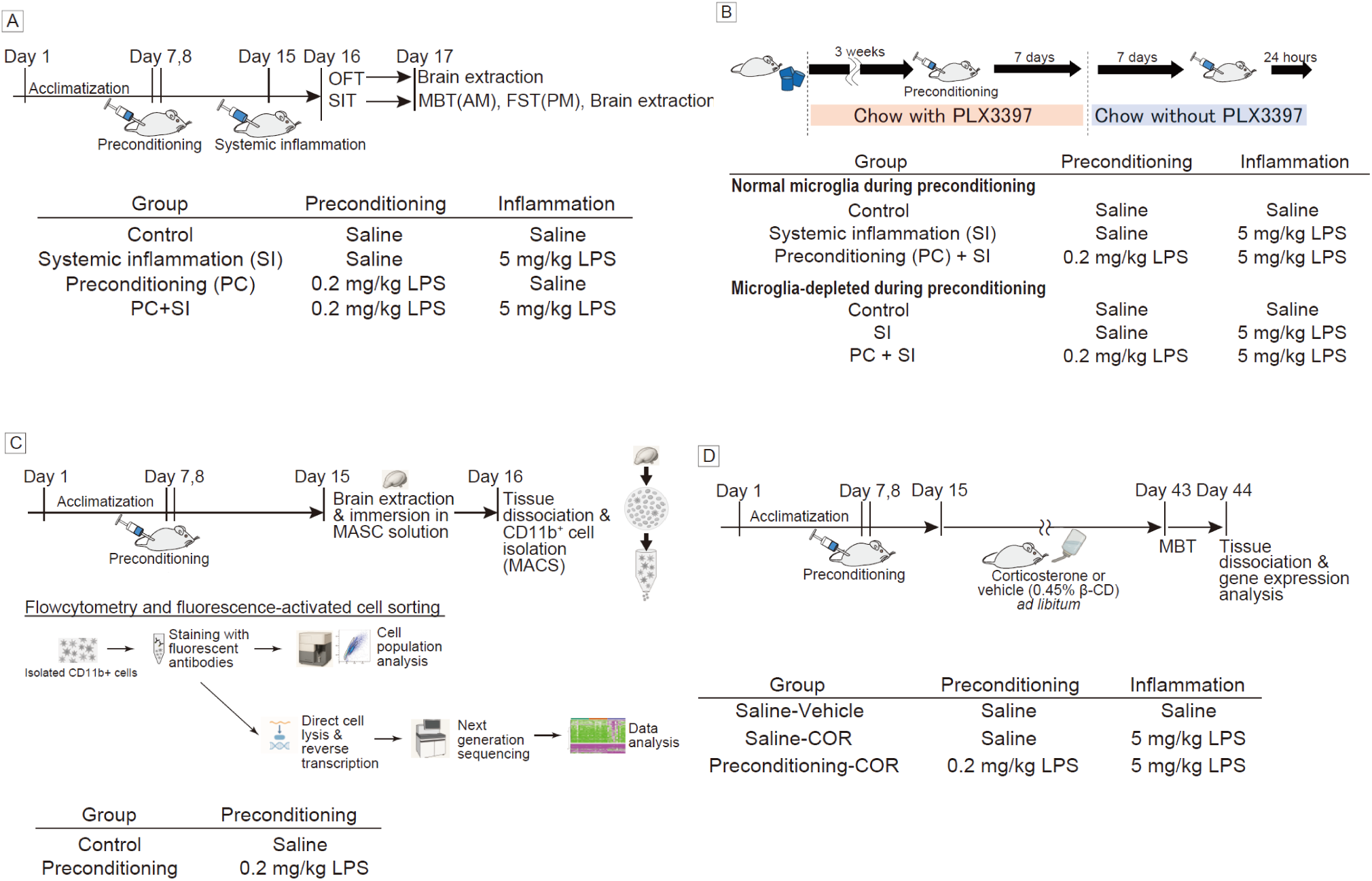
Experimental design and timeline. Schematic representation of three independent experiments. (A) Experiment 1: Effect of LPS preconditioning on behavioral abnormalities. Mice received saline or LPS (0.2 mg/kg) on days 1-2, followed by saline or LPS (5 mg/kg) on day 8. Behavioral tests were performed 24-48 hours post-challenge. (B) Experiment 2: Microglial depletion study. PLX3397 diet was administered for 3 weeks, followed by preconditioning and diet replacement to allow microglial repopulation before systemic inflammation challenge. (C) Experiment 3: Flow cytometry and RNA-seq analysis performed 7 days after preconditioning without systemic inflammation challenge. Illustration generated using ChatGPT (DALL·E), OpenAI.

### Experiment 1 (Figure 1A): Inhibitory effect of LPS preconditioning on behavioral abnormalities

Sixty-four mice were used across four independent replicates (16 mice per replicate). Mice were randomly assigned to four groups: (1) Control: saline on Days 1, 2, and 8; (2) Systemic inflammation (SI): saline on Days 1-2, LPS (5 mg/kg) on Day 8; (3) Preconditioning (PC): LPS (0.2 mg/kg) on Days 1-2, saline on Day 8; (4) PC+SI: LPS (0.2 mg/kg) on Days 1-2, LPS (5 mg/kg) on Day 8.

LPS (Escherichia coli O127:B8; Sigma-Aldrich, St. Louis, MO, USA) or saline (Otsuka Pharmaceutical, Naruto, Japan) was administered intraperitoneally at 10 mL/kg body weight. The preconditioning dose was based on previous neuroprotection studies (Rosenzweig et al., 2007), and the high-dose challenge was selected to induce neuroinflammation and behavioral changes (PMID: 31327964, 17203472). Behavioral tests were performed 24-48 hours post-challenge. Final sample sizes after attrition were: Control (n=12), SI (n=13), PC (n=11), PC+SI (n=11).

### Experiment 2 (Figure 1B): Verification of microglial necessity for behavioral protection

Forty-eight mice were divided into normal microglia (n=24) and microglia-depleted conditions (n=24). Microglial depletion was achieved using PLX3397-supplemented diet (290 mg/kg in AIN-76; CLEA Japan) for 3 weeks, as previously described ^29^. After dietary pretreatment, mice were subdivided into three groups per condition (initially n=8 each): Control (saline Days 1, 2, 16), SI (saline Days 1-2, LPS 5 mg/kg Day 16), and PC+SI (LPS 0.2 mg/kg Days 1-2, LPS 5 mg/kg Day 16). In microglia-depleted groups, PLX3397 diet was replaced with control diet on Day 9 to allow microglial repopulation. Final sample sizes: Groups 1-6: n=8, 6, 8, 8, 7, 6, respectively.

### Experiment 3 (Figure 1C): Brain macrophage subpopulation analysis

Twenty-four mice were randomly assigned to Control (n=12) or Preconditioning (n=12) groups. Preconditioning mice received LPS (0.2 mg/kg) on Days 1-2; controls received saline. Mice were sacrificed on Day 8 for flow cytometry and RNA-seq analysis. No attrition occurred in this experiment.

### Experiment 4 (Figure 1D): Effect of LPS preconditioning on chronic corticosterone administration model

To evaluate LPS preconditioning effects in a chronic stress model, six-week-old male C57BL/6N mice were purchased from Japan SLC (Hamamatsu, Japan) and randomly assigned to three groups (n=8 per group): Saline-Vehicle (control), Saline-COR (corticosterone exposure), and Preconditioning-COR (LPS preconditioning with corticosterone exposure). Mice received intraperitoneal injections of either low-dose LPS (0.2 mg/kg) or saline for two consecutive days (Days 1-2). Seven days after the final injection (Day 9), mice were provided with drinking water containing corticosterone (140 μg/mL; Tokyo Chemical Industry, Tokyo, Japan) dissolved in 0.45% β-cyclodextrin (β-CD; Tokyo Chemical Industry) for 4 weeks as previously described (Darcet et al., 2016). Control mice received vehicle (0.45% β-CD) only in their drinking water. Due to attrition during the experimental period, final sample sizes for behavioral analysis were: Saline-Vehicle (n=7), Saline-COR (n=6), and Preconditioning-COR (n=6).

### Analysis of brain immune cell composition changes induced by LPS preconditioning Biochemical analysis

After behavioral tests, the mice were anesthetized using an anesthetic mixture (0.75 mg/kg medetomidine [Medetomin injection, Meiji Seika Pharma Co., Ltd., Japan], 4.0 mg/kg midazolam [midazolam injection, TEVA, Takeda Pharmaceutical Co., Ltd., Japan], and 5 mg/kg butorphanol [Vetorphale, Meiji Seika Pharma Co., Ltd.]). The anesthetic mixture was injected at a volume of 10 mL/kg body weight. Then, cardio-perfusion with 0.1 M phosphate-buffered saline (PBS) was performed to remove blood. Next, the brains were extracted and submerged in 4% paraformalhehyde (PFA) and replaced the PFA with 30% sucrose in phosphate buffer and stored at 4 °C for immunohistochemistry.

### Open-field test

The OFT was conducted to assess spontaneous locomotor activity in a novel environment. The assessment was performed in an open-field arena, comprising an acrylic box 43.2 cm × 43.2 cm × 30.5 cm in size (Med Associates Inc., St. Albans, Vermont, USA). The sidewalls were covered with cardboard so that the mice could not see outside the open field. Horizontal and vertical arrays of 16 infrared beams tracked horizontal and vertical movements, respectively. Mice were placed in the center of the field and allowed to freely explore the chamber for 5 min in a dark room with a light source placed directly above the box. Feces and urine were removed from the arena and the arena was cleaned using 20% ethanol after each recording. This concentration was selected to avoid cracking of the acrylic apparatus, which can occur with higher ethanol concentrations (e.g., 70%). Activity Monitor version 5.50 software (Med associates Inc., Fairfax, Georgia, USA) was used to analyze the total distance moved by the mice.

### Sociability test

Sociability was evaluated using a modified open-field test, adapted from the protocol described by Lee et al. ^30^. The testing arena measured 40 × 30 × 30 cm and featured a central partition (20 × 35 × 1 cm) that divided the field into a U-shape configuration with two enclosed corners. In each corner, an acrylic enclosure (10.5 × 6.5 × 32 cm) with mesh-covered windows on three sides was placed, defining two regions of interest (ROI1 on the left and ROI2 on the right; each 15 × 12.5 cm). The enclosures were oriented so that the solid side faced the arena wall.

Test mice were acclimated to the experimental room for at least 30 minutes before testing. Initially, both enclosures were empty, and mice were allowed to freely explore the field for 5 minutes. After this habituation phase, the subject was returned to its home cage. A novel age-matched conspecific was then placed in the enclosure at ROI1. The test mouse was reintroduced and allowed to explore for another 5 minutes. All sessions were video recorded, and the time spent in each ROI was quantified using TimeOFCR1 software (O’Hara & Co., Tokyo, Japan). A preference score was calculated as the difference in time spent between ROI1 and ROI2 divided by the total session time (300 seconds), with a score of 0 indicating no preference.

### Forced swim test

The forced swim test (FST), commonly used to assess depressive-like behavior in rodents, was performed based on a previously described protocol^31^, with the addition of an automated behavioral analysis system, MicroAct™ (Neuroscience, Japan) ^32^.

Immediately before the test, a small magnet (1.6 mm in diameter and 0.5 mm thick) was attached to the heel of each hind paw using instant adhesive. Mice were then placed individually into a transparent acrylic cylinder (18 cm height × 11 cm diameter) filled with water (maintained at 22–23 °C) to a depth of 12.5 cm. The cylinder was surrounded by coils that detected movements of the attached magnets via electromagnetic induction. The induced current was amplified, converted to voltage, and recorded automatically.

Immobility time was measured during the final 5 minutes of the 6-minute test session. The system was configured to detect voltage pulses within a frequency range of 1–30 Hz and an amplitude threshold above 0.10 V. Immobility was defined as the total duration (in seconds) of all intervals exceeding 1.00 s between consecutive voltage pulses, reflecting minimal or absent movement.

### Marble burying test

The marble burying test, widely used to assess anxiety-like behavior, was performed based on previously reported methods (Kedia et al., 2014). Mice exhibiting anxiety-like behavior bury a greater number of marbles in this test. A clean cage (225 × 338 × 140 mm) was filled with sawdust bedding to a depth of 5 cm. For habituation, mice were individually placed in the cage for 30 minutes. Mice were then removed, the bedding was leveled, and 15 glass marbles were arranged in a 3 × 5 grid pattern at equal intervals on the surface. Mice were then returned to the same cage for 30 minutes. The number of marbles buried or partially buried (≥ two-thirds covered by bedding) was counted.

### Immunohistochemistry

Mice were deeply anesthetized using an anesthetic mixture (0.75 mg/kg medetomidine [Medetomin injection, Meiji Seika Pharma Co., Ltd., Japan], 4.0 mg/kg midazolam [midazolam injection, TEVA, Takeda Pharmaceutical Co., Ltd., Japan], and 5 mg/kg butorphanol [Vetorphale, Meiji Seika Pharma Co., Ltd.]). The anesthetic mixture was injected at a volume of 10 mL/kg. Then, cardio-perfusion with 0.1 M phosphate buffer (PB) was performed to remove the blood. Next, the brains were extracted and immersed in 4% paraformaldehyde in 0.1M phosphate buffer for fixation. In the next day, the paraformaldehyde solution was removed and immersed the brain with 30% sucrose in PB. The brains in the buffer were kept at 4°C until making slices. Coronal brain sections (30 μm thickness) were prepared using a cryostat (CryoStar NX70, Epredia, USA). Sections were stored at 4 °C in phosphate buffer (PB) containing 0.05% sodium azide until staining. For antigen retrieval, sections at approximately Bregma –2.13 mm were mounted on glass slides (SCRE, Matsunami, Osaka, Japan) and air-dried overnight. Slides were then immersed in 10 mM sodium citrate buffer (pH 6.0) preheated to 98 °C and incubated at this temperature for 30 min using a glass jar. Subsequently, the jar was left at room temperature for 30 min to allow gradual cooling.

Slides were then rinsed in ultrapure water for 15 min and permeabilized with 0.2% Triton X-100 in PB (PBTx) for 15 min. After rinsing with 0.1 M phosphate buffer to remove detergent, sections were incubated in blocking solution (10% normal donkey serum [NDS] in PBTx) for 1 h at room temperature. Primary antibodies diluted in 3% NDS in PBTx were applied, and the sections were incubated overnight at 4 °C in a humidified chamber. The following primary antibodies were used: anti-IBA1 (1:500, Cat# 234006, Synaptic Systems, Göttingen, Germany), anti-GFAP (1:1000, Cat# ab3554, Abcam, Cambridge, UK), and anti-PBR (1:200, Cat# ab109497, Abcam).

After washing with PBTx, the sections were incubated with secondary antibodies for 2 h at room temperature in the dark. Secondary antibodies included Alexa Fluor 488-conjugated donkey anti-chicken IgY (1:400, Cat# 703-545-155, Jackson ImmunoResearch, West Grove, PA, USA), Alexa Fluor 568-conjugated donkey anti-goat IgG (1:400, Cat# A32849, Thermo Fisher Scientific, Waltham, MA, USA), and Alexa Fluor 647-conjugated donkey anti-rabbit IgG (1:400, Cat# A10042, Thermo Fisher Scientific). Nuclear staining was performed with DAPI (0.1 μg/mL) for 10 min at room temperature in the dark, followed by washing with PBTx.

Sections were mounted using ProLong™ Gold Antifade Mountant (Thermo Fisher Scientific) and covered with coverslips. After drying for 24 h at room temperature in the dark, images were acquired using a fluorescence microscope (BZ-X710, Keyence, Osaka, Japan) with a 20× objective lens in sectioning mode.

Signal area analysis of PBR, IBA1, and GFAP was performed using the hybrid cell count function of Keyence BZ-X800 Analyzer (Keyence). Images of 200×300 μm regions from hippocampal CA1 and CA3 areas were cropped from the microscopic images and used for analysis. In this study, intensity normalization was performed for each fluorescence channel using histogram stretching. Specifically, pixel intensity values were linearly transformed based on the detected minimum and maximum values, and rescaled to span the full 16-bit dynamic range (0–65535). This procedure corrected for inter-sample variability in image brightness and ensured consistency and validity in subsequent inter-channel quantitative comparisons. Following normalization, the images were merged into overlay composites, from which spatially colocalized regions were extracted and the co-expression area was quantitatively and objectively measured.

### Brain macrophages isolation and CD11b positive cell collection

Whole mouse brains were sagittally sectioned into eight pieces using a sterile razor blade. The tissue fragments were placed on a stainless-steel mesh (pore size: 77 μm, 0.05 mm wire diameter; Tokyo Screen Co., Ltd., Japan) set in a 100 mm dish containing 20 mL of HBSS (with calcium and magnesium, supplemented with 20 mM HEPES, pH 7.2; Thermo Fisher Scientific). Gentle dissociation was performed using the plunger of a syringe. The resulting suspension was filtered through a 70 μm pluriStrainer (PLS, Germany) into a 50 mL conical tube. An additional 20–25 mL of HBSS was applied to the mesh to collect residual cells, and the combined suspension (∼40 mL) was centrifuged at 300 × g for 5 min at 4°C. The supernatant was removed, and the cell pellet was resuspended in 5.2 mL of HBSS.

For debris removal, 1.8 mL of pre-chilled Debris Removal Solution (Miltenyi Biotec) was added and mixed gently. Then, 4 mL of HBSS was carefully layered on top to form a gradient, followed by centrifugation at 3000 × g for 10 min at 4°C. The top two layers were aspirated, and the remaining fraction was brought up to 15 mL with HBSS, gently mixed, and centrifuged at 1000 × g for 10 min at 4°C. The pellet was resuspended in 1× Red Blood Cell Removal Solution (Miltenyi Biotec), incubated for 10 min at 4°C, and quenched with 10 mL of FACS buffer (PBS without calcium and magnesium, supplemented with 0.5% BSA and 15 mM HEPES, pH 7.2). After centrifugation at 300 × g for 5 min at 4°C, the pellet was resuspended in 100 μL of FACS buffer.

20 μL of CD11b (Microglia) MicroBeads, human and mouse (Miltenyi Biotec) was added and incubated on ice for 15 min. Following incubation, 1 mL of FACS buffer per 1×10^7^ cells was added and gently mixed, then centrifuged at 300 × g for 5 min. The pellet was resuspended in 500 μL of FACS buffer per 1×10^7^ cells and subjected to magnetic separation using LS columns (Miltenyi Biotec) according to the manufacturer’s instructions. Columns were equilibrated with FACS buffer, and labeled cells were applied. The columns were washed three times with 300μL of FACS buffer. CD11b^+^ cells were eluted by removing the column from the magnetic separator and flushing with 5 mL of FACS buffer using a plunger, followed by an additional rinse. The collected CD11b^+^ cell suspension was centrifuged at 300 × g for 5 min, and the pellet was used for downstream analysis.

### Flow cytometry

Collected CD11b^+^ cells were incubated with an Fc-blocker (BD Biosciences, Franklin Lakes, NJ, USA) to prevent non-specific binding. For the analysis of CNS macrophage subpopulations based on TMEM119 and CD45 expression, CD11b-positive cells were incubated on ice for 15 min with PE-conjugated anti-TMEM119 (V3RT1GOsz, eBioscience, San Diego, CA, USA) and BV421-conjugated anti-CD45 (30-F11, eBioscience). Following incubation, cells were washed twice with phosphate-buffered saline containing 0.5% bovine serum albumin and 10 mM HEPES at pH 7.4. For live-dead analysis, cells were stained with Zombie NIR viability kit (BioLegend) before the incubation with an Fc-blocker and fluorescent antibody staining.

Flow cytometry was performed using a BD FACSCanto II instrument, and data were analyzed using FlowJo software (BD Biosciences). Forward scatter (FSC-A) and side scatter (SSC-A) were used as indicators of cell size and internal complexity, respectively. FSC-H was additionally employed to discriminate single cells from doublets using FSC-A vs. FSC-H plots.

The analysis was conducted using cells isolated from the brains of 12 control mice and 12 mice preconditioned with low-dose lipopolysaccharide (Pre-LPS group).

### Cell sorting

Collected CD11b+ cells were incubated with an Fc-blocker (BD Biosciences, Franklin Lakes, NJ, USA) to prevent non-specific binding. For the analysis of CNS macrophage subpopulations based on TMEM119 and CD45 expression, CD11b-positive cells were incubated on ice for 15 min with PE-conjugated anti-TMEM119 (V3RT1GOsz, eBioscience) and APC-conjugated anti-CD45 (30-F11, eBioscience). Following incubation, cells were washed twice with phosphate-buffered saline containing 0.5% bovine serum albumin and 10 mM HEPES at pH 7.4. Cell sorting was performed using a SH800S instrument (Sony, Japan). Forward scatter (FSC-A) and side scatter (SSC-A) were used as indicators of cell size and internal complexity, respectively. FSC-H was additionally employed to discriminate single cells from doublets using FSC-A vs. FSC-H plots. The analysis was conducted using cells isolated from the brains of 12 control mice and 12 mice preconditioned with low-dose lipopolysaccharide (Pre-LPS group).

### RNA extraction and quantitative PCR analysis

For gene expression analysis in the corticosterone model, mice were deeply anesthetized with an anesthetic mixture (0.75 mg/kg medetomidine hydrochloride, 4.0 mg/kg midazolam, and 5.0 mg/kg butorphanol tartrate) administered intraperitoneally, and transcardially perfused with sterile PBS. One hemisphere was immersed in RNAlater solution (Thermo Fisher Scientific) and stored at 4°C. The hippocampus was dissected and stored at -80°C until RNA extraction.

Total RNA was extracted from hippocampal tissue using the RNeasy Lipid Tissue Mini Kit (Qiagen). Complementary DNA was synthesized from 2 μg of total RNA using ReverTra Ace qPCR RT Master Mix (Toyobo). The cDNA was diluted 1:20 in 2.5 M betaine (Sigma-Aldrich) aqueous solution. Quantitative real-time PCR was performed in triplicate using Thunderbird SYBR qPCR Mix (Toyobo) on a LightCycler 480 system (Roche).

Amplification parameters were: initial denaturation at 95°C for 1 min, followed by 50 cycles of 95°C for 15 s and 60°C for 1 min. Target genes included *Il1b* (inflammatory cytokine), *Tlr4* (Toll-like receptor 4), and *Myd88* (myeloid differentiation factor 88), with *Actb* (β-actin) as the internal control. Primer sequences are provided in Table 1. Gene expression levels were compared among experimental groups using the 2^-ΔΔCt^ method (Livak and Schmittgen, 2001). Outliers were identified and excluded using the interquartile range method. Final sample sizes for gene expression analysis were: *Il1b*: Saline-Vehicle (n=7), Saline-COR (n=6), LPS-COR (n=5); *Tlr4*, *Myd88*: Saline-Vehicle (n=6), Saline-COR (n=6), LPS-COR (n=6).

**Table 1.**
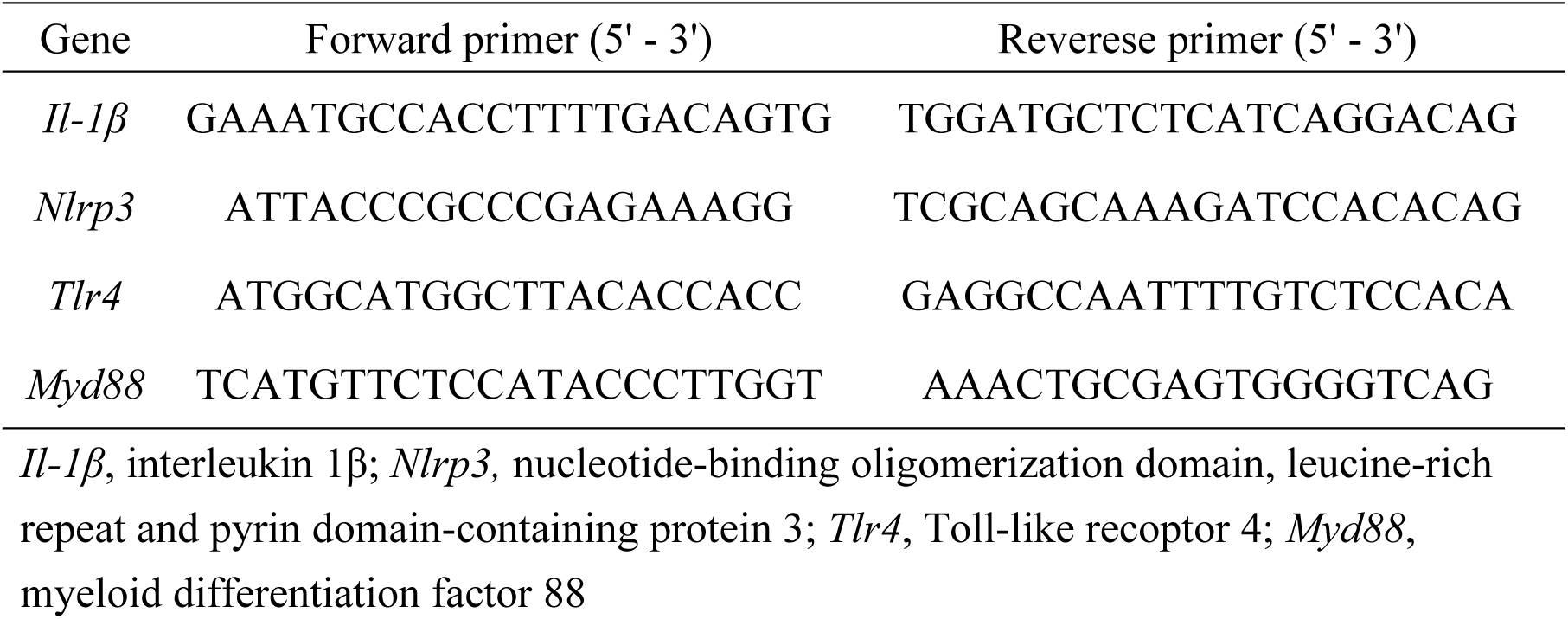
Primers for quantitative PCR.

### Bulk RNA sequencing analysis

Target brain macrophage subpopulations were isolated by fluorescence-activated cell sorting (FACS) as described above. Due to the low cell number (<1,000 cells per sample), RNA was not extracted; instead, full-length cDNA synthesis was directly performed from cell lysates using the SMART-Seq mRNA kit (Takara Bio) according to the manufacturer’s low-input protocol. Libraries were prepared using the Nextera XT DNA Library Prep Kit (Illumina), involving tagmentation and barcode indexing, and their quality was assessed using the Agilent TapeStation system (High Sensitivity D1000).

Sequencing was performed on an Illumina platform with paired-end 150-bp reads (2 × 150 bp), yielding approximately 10–60 million read pairs per sample. Reads were processed and mapped to the mouse reference genome (GRCm39, GENCODE release M28) using DRAGEN Bio-IT Platform (v4.3.6). Gene- and transcript-level expression values were calculated as transcripts per million (TPM) and raw read counts.

Differential gene expression analysis was performed using edgeR (v3.40.2). Genes were filtered using the filterByExpr function, and exact tests were applied to compare groups. Genes with fold change >1 and p-value < 0.05 were considered upregulated, and those with fold change <1 and p-value < 0.05 were considered downregulated.

Gene enrichment analysis was conducted using gprofiler2 (v0.2.1) and clusterProfiler (v4.6.2), referencing Gene Ontology (Biological Process, Molecular Function, Cellular Component), Reactome, and WikiPathways databases. Multiple testing correction was performed using the Benjamini–Hochberg method. When applicable, motif enrichment analysis was also conducted using RcisTarget (v1.18.2) to identify overrepresented transcription factor binding motifs and putative upstream regulators.

### Statistical analysis

Statistical analyses were performed using GraphPad Prism 9.5.1 (GraphPad Software, La Jolla, CA, USA). Data are expressed as mean ± SEM. For behavioral and immunohistochemical data, two-way ANOVA followed by Tukey’s post-hoc test was used for multiple comparisons. For flow cytometry data comparing two groups, unpaired Student’s t-test was applied. For comparisons among multiple groups in microglial depletion experiments, one-way ANOVA with Tukey’s post-hoc test was performed.

RNA-sequencing data were analyzed as described in the Bulk RNA sequencing analysis section. Statistical significance was set at p < 0.05, with p-values between 0.05 and 0.1 considered as trends.

Animals requiring euthanasia due to severe illness or injury were excluded from final analysis according to predetermined criteria. Sample attrition was primarily due to fighting-related injuries and mortality following high-dose LPS administration, which is an inherent limitation of this model.

## Result

### 1. LPS preconditioning attenuates systemic inflammation-induced behavioral changes

To evaluate the effects of low-dose LPS preconditioning on behavioral changes following systemic inflammation, we performed open-field test, forced swim test, and social interaction test.

Regarding the open-field test, two-way ANOVA of total distance traveled revealed significant main effects of systemic inflammation (F(1,43) = 48.30, p < 0.0001) and preconditioning (F(1,43) = 6.098, p < 0.05), with no significant interaction (F(1,43) = 2.616, p = 0.113). Analysis of main effects indicated that systemic inflammation decreased total distance traveled, while preconditioning tended to increase locomotor activity regardless of inflammation status (Figure 2A).

**Figure 2.**
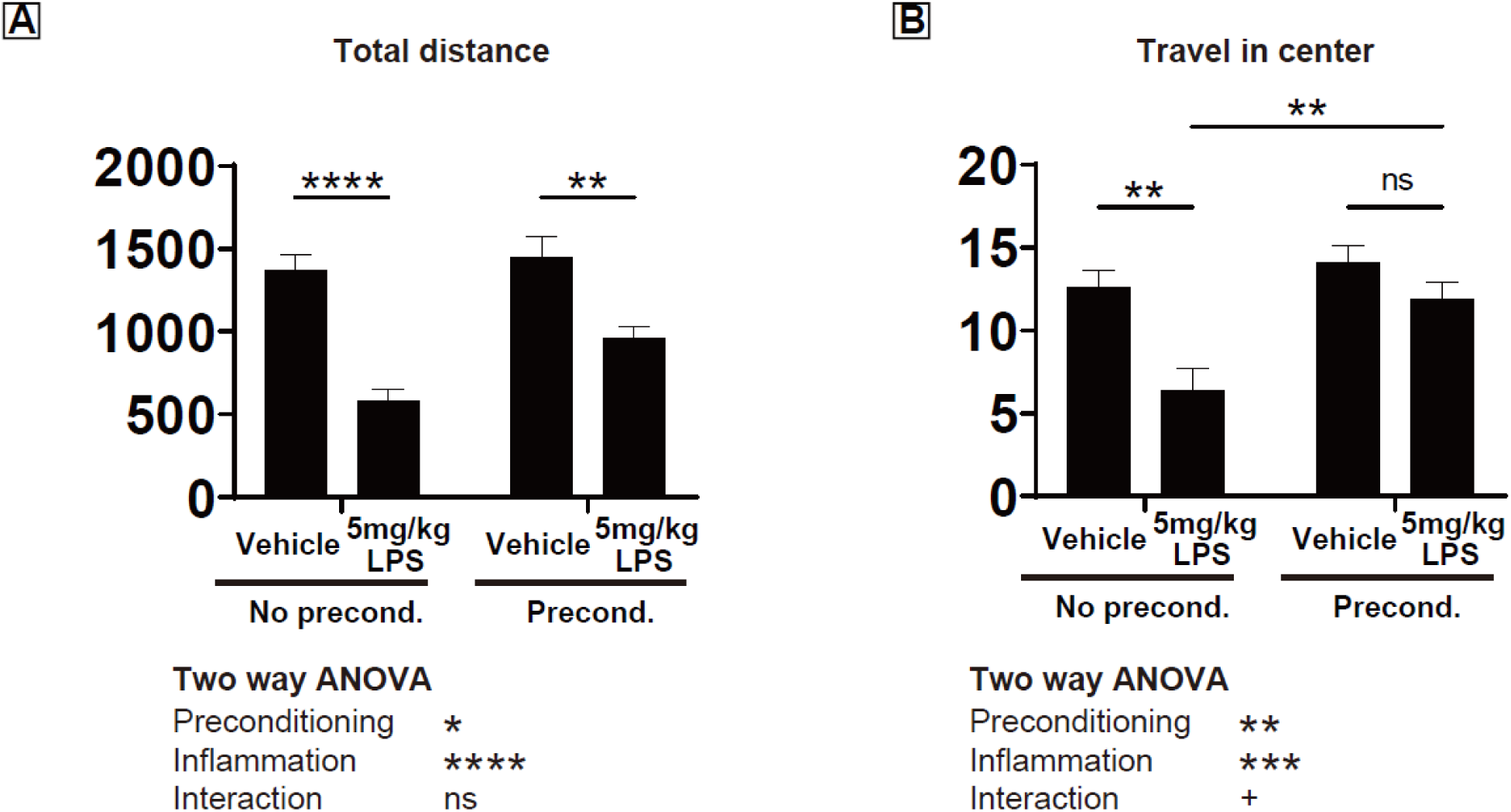
LPS preconditioning effects on locomotor activity. (A) Total distance traveled in open field test. (B) Distance traveled in center zone. Two-way ANOVA with Tukey’s post-hoc test. n = 11-13 per group. *p < 0.05, **p < 0.01. Data represent mean ± SEM.

For center distance, two-way ANOVA showed significant main effects of systemic inflammation (F(1,43) = 13.60, p < 0.001) and preconditioning (F(1,43) = 9.510, p < 0.01). The interaction approached significance (F(1,43) = 3.195, p = 0.081). Exploratory post-hoc Tukey’s test revealed that in non-preconditioned mice, systemic inflammation significantly reduced center distance (No precond.-Vehicle: 12.6 ± 1.0 cm vs. No precond.-5mg/kg LPS: 6.4 ± 1.4 cm, p < 0.01), whereas preconditioned mice showed no significant reduction following systemic inflammation (Precond.-Vehicle: 14.1 ± 1.0 cm vs. Precond.-5mg/kg LPS: 11.9 ± 1.0 cm, p > 0.05), suggesting preconditioning may attenuate anxiety-like behavior induced by systemic inflammation (Figure 2B).

Regarding the forced swim test, two-way ANOVA of immobility time revealed a significant main effect of systemic inflammation (F(1,55) = 4.052, p < 0.05), but no main effect of preconditioning (F(1,55) = 1.422, p > 0.05) or interaction (F(1,55) = 2.490, p > 0.05). These results indicate that while systemic inflammation tended to increase immobility time, preconditioning showed no clear protective effect on depression-like behavior (Figure 3).

**Figure 3.**
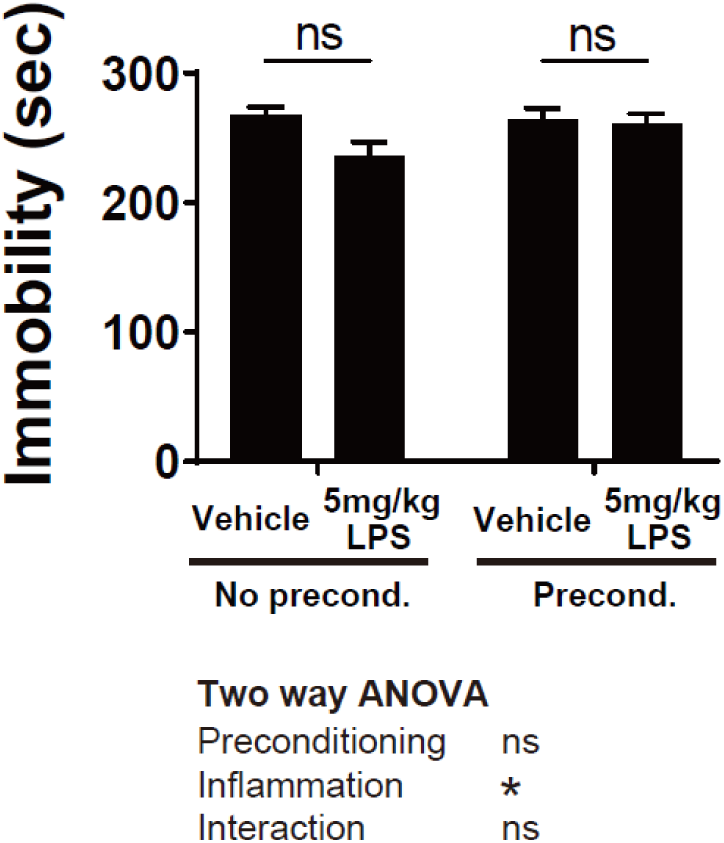
Forced swim test. Immobility time during the last 5 minutes of 6-minute test session. Two-way ANOVA. n = 11-13 per group. Data represent mean ± SEM.

To assess social behavior, we performed the U-shaped social interaction test. Initially, exploratory behavior was evaluated with both ROIs empty, confirming no significant differences in time spent between ROI1 and ROI2 across all groups, indicating absence of place preference bias (Figure 4A).

**Figure 4.**
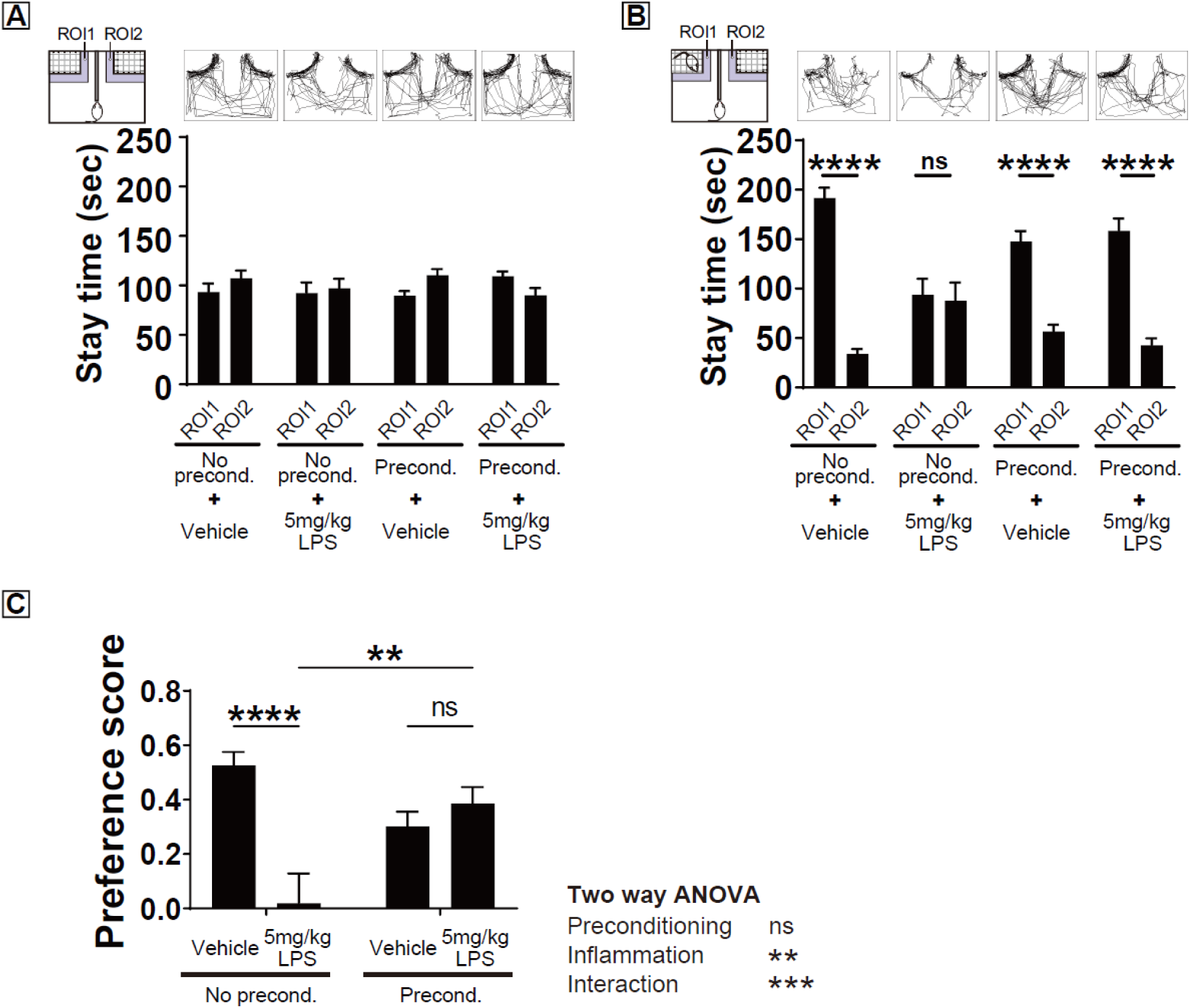
Social interaction behavior. (A) Time spent in ROI1 and ROI2 with both enclosures empty. (B) Time spent in ROI1 (with conspecific) and ROI2 (empty) for each group. (C) Social preference score calculated as (ROI1-ROI2)/300 seconds. Two-way ANOVA with Tukey’s post-hoc test. n = 11-13 per group. **p < 0.01, ****p < 0.0001. Data represent mean ± SEM.

When an age-matched male mouse was placed in ROI1, control mice (No precond.-Vehicle) spent significantly more time in ROI1 compared to empty ROI2 (191.5 ± 10.5 s vs. 33.9 ± 5.0 s, p < 0.0001). In contrast, mice with systemic inflammation (No precond.-5mg/kg LPS) showed no significant preference (93.6 ± 16.3 s vs. 87.6 ± 18.6 s, p > 0.05). Preconditioned mice maintained significant preference for ROI1 in both vehicle (147.3 ± 11.0 s vs. 56.8 ± 6.7 s, p < 0.0001) and LPS-treated groups (158.5 ± 12.5 s vs. 42.8 ± 6.9 s, p < 0.0001) (Figure 4B).

Two-way ANOVA of social preference scores revealed a significant main effect of systemic inflammation (F(1,55) = 8.887, p < 0.01) and a significant interaction (F(1,55) = 17.37, p < 0.001), with no main effect of preconditioning (F(1,55) = 1.012, p > 0.05). Post-hoc analysis showed that systemic inflammation significantly reduced social preference scores in non-preconditioned mice (No precond.-Vehicle: 0.53 ± 0.05 vs. No precond.-5mg/kg LPS: 0.02 ± 0.11, p < 0.0001), while preconditioned mice showed no such reduction (Precond.-Vehicle: 0.30 ± 0.05 vs. Precond.-5mg/kg LPS: 0.39 ± 0.06, p > 0.05). Moreover, under systemic inflammation, preconditioned mice showed significantly higher social preference scores compared to non-preconditioned mice (No precond.- 5mg/kg LPS: 0.02 ± 0.11 vs. Precond.-5mg/kg LPS: 0.39 ± 0.06, p < 0.01) (Figure 4C).

### 2. Effects of LPS preconditioning are accompanied by changes in brain glial cells

To investigate the neural basis of behavioral changes, we performed immunohistochemical analysis of neuroinflammation-related markers in hippocampal CA1 and CA3 regions. We quantitatively evaluated expression of microglial marker (IBA1), astrocyte marker (GFAP), and glial activation marker (PBR/TSPO) (Figure 5A).

**Figure 5.**
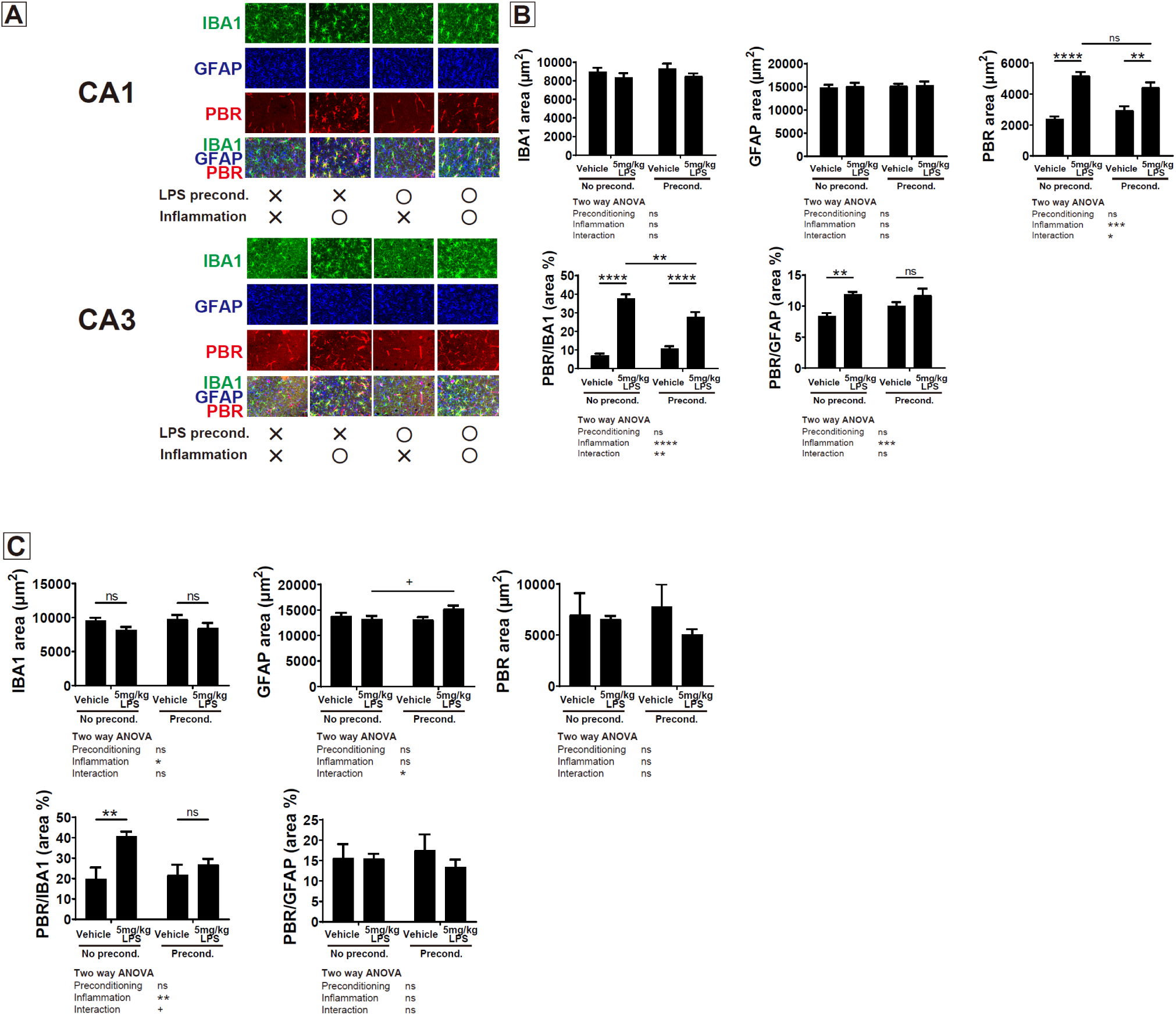
Immunohistochemical analysis of neuroinflammation markers in hippocampus. (A) Representative images of IBA1, GFAP, and PBR immunostaining in CA1 and CA3 regions. Scale bar = 50 μm. (B) Quantification of signal areas in CA1 region. (C) Quantification of signal areas in CA3 region. Two-way ANOVA with Tukey’s post-hoc test. n = 7-8 per group. *p < 0.05, **p < 0.01, ****p < 0.0001. Data represent mean ± SEM.

#### Changes in CA1 region

In CA1 region, two-way ANOVA of IBA1 and GFAP signal areas revealed no significant main effects of preconditioning, systemic inflammation, or their interaction (Figure 5B).

For PBR (peripheral benzodiazepine receptor) signal area, two-way ANOVA showed significant main effect of systemic inflammation (F(1,26) = 77.00, p < 0.0001) and interaction (F(1,26) = 7.349, p < 0.05), with no main effect of preconditioning (F(1,26) = 7.349, p > 0.05). Post-hoc Tukey’s test revealed that systemic inflammation increased PBR signal in both conditions (No precond.-Vehicle: 2368.0 ± 185.6 μm² vs. No precond.-5mg/kg LPS: 5194.9 ± 233.3 μm², p < 0.0001; Precond.-Vehicle: 2953.4 ± 261.4 μm² vs. Precond.-5mg/kg LPS: 4445.9 ± 305.2 μm², p < 0.01), with no significant difference between inflamed groups.

For PBR signal area overlapping with IBA1-positive regions, two-way ANOVA revealed significant main effect of systemic inflammation (F(1,26) = 144.9, p < 0.0001) and interaction (F(1,26) = 12.14, p < 0.01). Post-hoc analysis showed that systemic inflammation increased PBR/IBA1 ratio in both conditions (No precond.-Vehicle: 7.0 ± 0.9% vs. No precond.-5mg/kg LPS: 37.5 ± 2.4%, p < 0.0001; Precond.-Vehicle: 10.8 ± 1.2% vs. Precond.-5mg/kg LPS: 27.6 ± 2.8%, p < 0.0001), but preconditioning significantly suppressed this increase (No precond.-5mg/kg LPS: 37.5 ± 2.4% vs. Precond.-5mg/kg LPS: 27.6 ± 2.8%, p < 0.01).

For PBR signal area overlapping with GFAP-positive regions, only the main effect of systemic inflammation was significant (F(1,26) = 14.07, p < 0.001).

#### Changes in CA3 region

In CA3 region, two-way ANOVA revealed a significant main effect of systemic inflammation on IBA1 signal area (F(1,26) = 5.308, p < 0.05) (Figure 5C). For GFAP signal area, the interaction was significant (F(1,26) = 5.576, p < 0.05), but post-hoc testing showed only a trend between inflamed groups (No precond.-5mg/kg LPS: 13119.8 ± 652.3 μm² vs. Precond.-5mg/kg LPS: 15295.1 ± 596.7 μm², p < 0.1). PBR signal area showed no significant main effects or interaction.

For PBR/IBA1 ratio, two-way ANOVA revealed a significant main effect of systemic inflammation (F(1,26) = 9.459, p < 0.01) and a trend for interaction (F(1,26) = 3.473, p < 0.1). Post-hoc testing showed that only non-preconditioned mice had significant increase in PBR/IBA1 ratio with systemic inflammation (No precond.-Vehicle: 19.7 ± 5.7% vs. No precond.-5mg/kg LPS: 40.7 ± 2.4%, p < 0.01). PBR signal area overlapping with GFAP-positive regions showed no significant effects.

### 3. Microglia are required to induce protective effect of LPS preconditioning

To verify the necessity of microglia for the protective effect of preconditioning, we performed temporally controlled microglial depletion experiments. Administration of CSF1R inhibitor PLX3397-containing diet for 3 weeks resulted in marked reduction of IBA1-positive cells, consistent with the report by Elmore et al. (2014) (Figure 6, upper panel).

**Figure 6.**
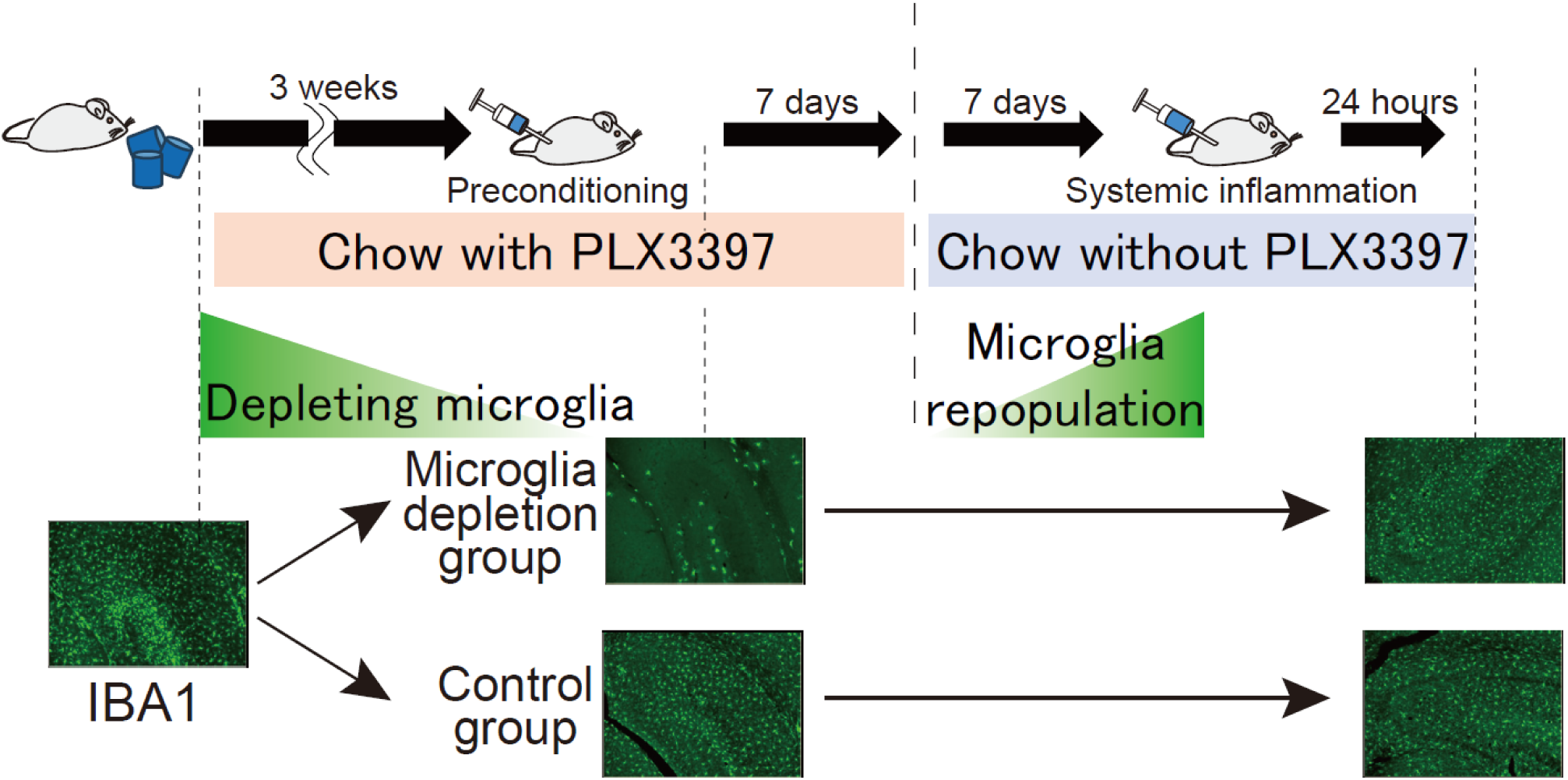
Microglial depletion and repopulation. Representative images of IBA1 immunostaining showing microglial depletion after 3 weeks of PLX3397 diet (upper panels) and repopulation after diet replacement (lower panels). Scale bar = 100 μm.

In this study, we employed the following experimental design to specifically deplete microglia during the preconditioning period: After 3 weeks of PLX3397 administration (when microglia were depleted), low-dose LPS (0.2 mg/kg) preconditioning was performed, and PLX3397-containing diet was replaced with control diet 7 days later. This 7-day period was based on the time required for preconditioning effects to establish in our study. Following an additional 7-day recovery period after diet replacement, IBA1-positive cell numbers recovered through proliferation of residual microglia (Figure 6, lower panel). This experimental system created conditions where microglia were depleted only during preconditioning induction, but were present during subsequent systemic inflammation induction.

Using the U-shaped social interaction test, we evaluated social preference scores calculated from time spent in ROI1 (with conspecific) versus empty ROI2. One-way ANOVA revealed significant differences among groups (F(5,37) = 5.680, p < 0.001). Post-hoc testing showed that in mice with normal microglia, systemic inflammation significantly reduced social preference scores (Control: 0.27 ± 0.05 vs. Inflammation: -0.49 ± 0.19, p < 0.001), and preconditioning significantly suppressed this reduction (Inflammation: -0.49 ± 0.19 vs. Precond.-Inflammation: 0.14 ± 0.10, p < 0.01).

In contrast, in mice with microglia depleted during preconditioning, systemic inflammation reduced social preference scores (Control: 0.31 ± 0.09 vs. Inflammation: -0.15 ± 0.16, p < 0.05), but preconditioning failed to improve social preference scores (Inflammation: -0.15 ± 0.16 vs. Precond.- Inflammation: -0.09 ± 0.15, p > 0.05) (Figure 7).

**Figure 7.**
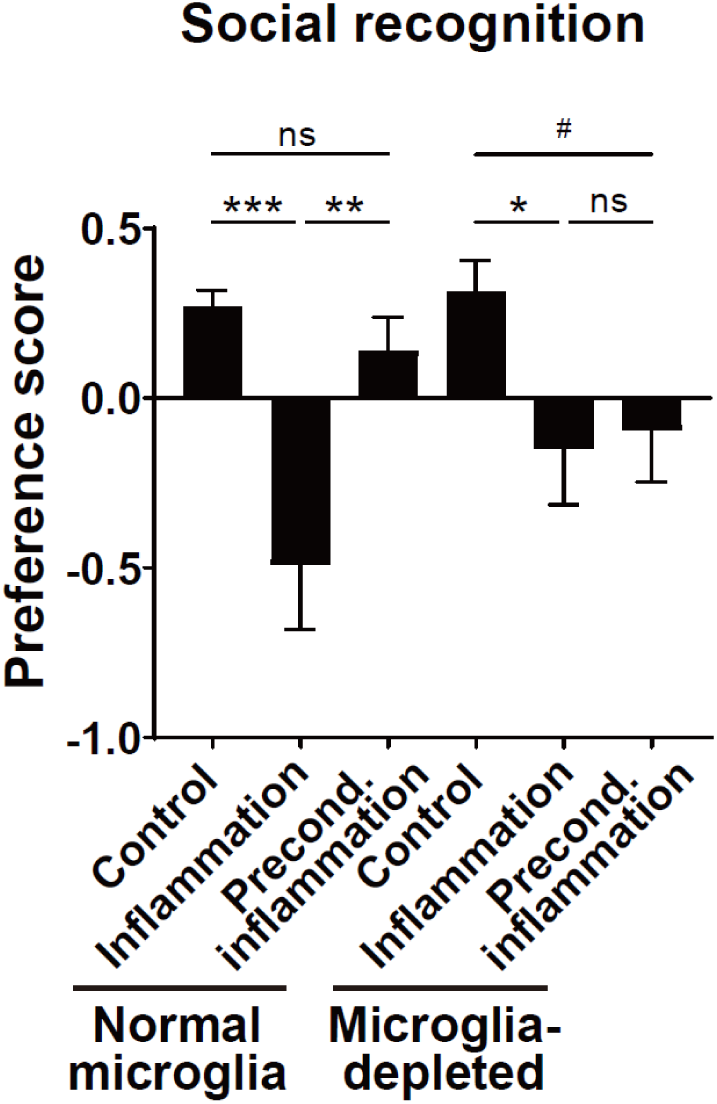
Microglial requirement for preconditioning effects. Social preference scores in mice with normal or depleted microglia during preconditioning period. One-way ANOVA with Tukey’s post-hoc test. n = 6-8 per group. *p < 0.05, **p < 0.01, ***p < 0.001. Data represent mean ± SEM.

### 4. LPS preconditioning alters phenotypes of brain macrophages

To investigate the effects of preconditioning on brain macrophage composition, we performed flow cytometric analysis. CD11b-positive cells were isolated from brains of Control mice (no preconditioning, n=11) and Preconditioning mice (n=11), and analyzed for TMEM119 and CD45 expression patterns.

After excluding debris using forward scatter (FSC) and side scatter (SSC) (Figure 8A), CD45-positive cells were gated (Figure 8B). Using FSC-A vs FSC-H plots to exclude doublets from CD45-positive cells, we identified two distinct populations: Single cells 1 and Single cells 2 (Figure 8C). Additionally, a TMEM119-negative CD45-positive cell population was observed (Figure 8D). For exclusion of dead cells and cell doublets, we performed a viability test and doublet cell exclusion was carefully performed using intensive analysis of FSC and SSC fluorescent signals (Supplementary Fig. 2).

**Figure 8.**
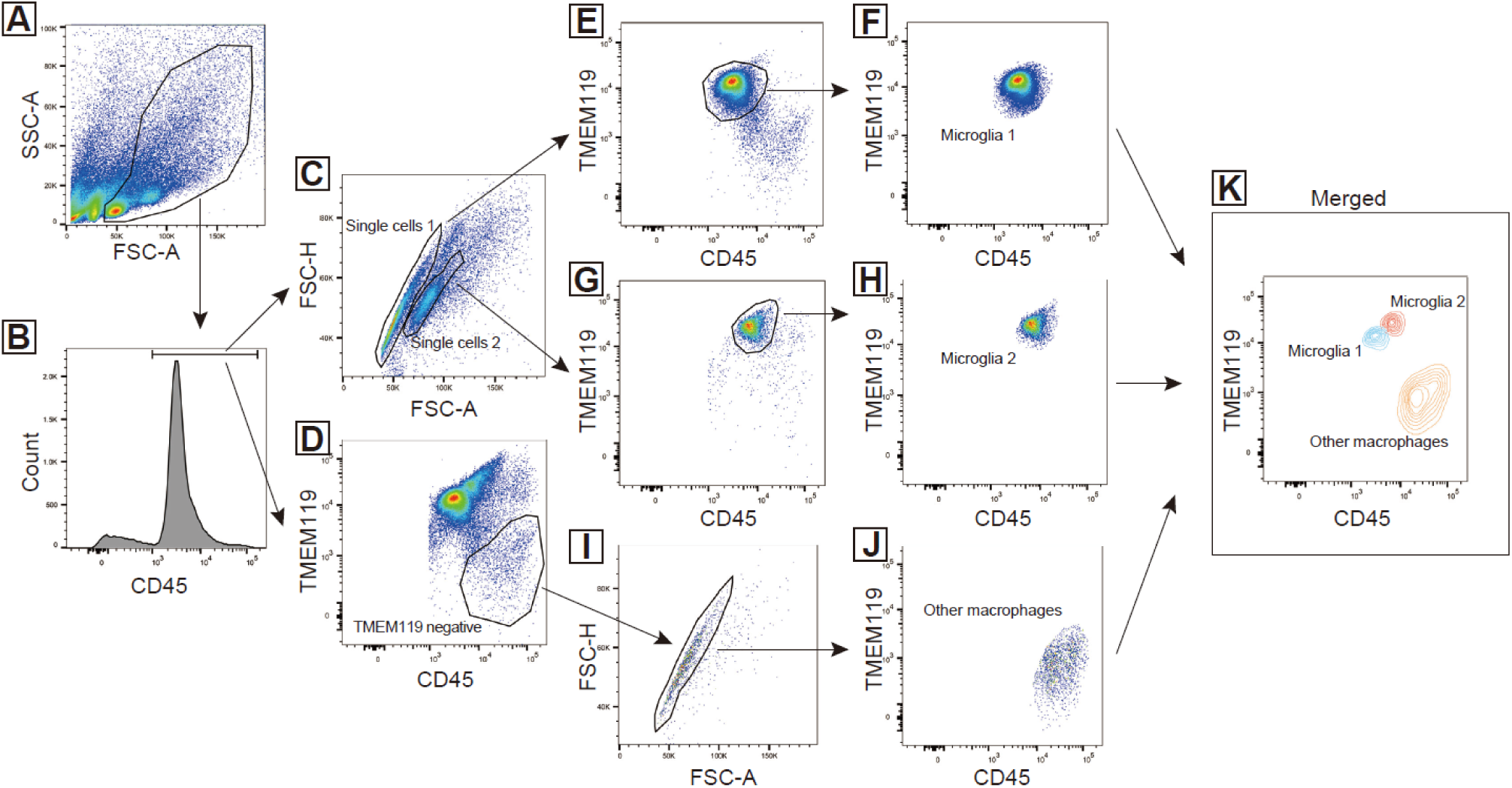
Flow cytometric identification of brain macrophage subpopulations. (A-J) Sequential gating strategy for identifying brain macrophage subpopulations. (A) Debris exclusion. (B) CD45+ cell selection. (C) Doublet exclusion. (D) TMEM119-negative population. (E-H) Identification of Microglia 1 and Microglia 2. (I-J) Identification of Other brain macrophages. (K) Overlay plot showing three distinct subpopulations based on TMEM119 and CD45 expression.

From Single cells 1 and Single cells 2, after excluding doublets and TMEM119-negative cells, we identified Microglia 1 (TMEM119-positive CD45-low) and Microglia 2 (TMEM119-positive CD45-intermediate), respectively (Figure 8E-H). From TMEM119-negative cells, after doublet exclusion, we identified other brain macrophages (TMEM119-negative CD45-high) (Figure 8I, J). These three subpopulations were shown in an integrated plot based on TMEM119 and CD45 expression patterns (Figure 8K).

Comparison of subpopulation proportions among CD45-positive cells revealed that Pre-LPS mice showed significantly decreased Microglia 1 proportion compared to Control (Control: 0.82 ± 0.015% vs. Pre-LPS: 0.76 ± 0.015%, t(20) = 2.902, p < 0.01) (Figure 9A), and significantly increased Microglia 2 proportion (Control: 0.072 ± 0.008% vs. Pre-LPS: 0.11 ± 0.007%, t(20) = 3.946, p < 0.001) (Figure 9B). No significant change was observed in Other brain macrophages proportion (Control: 0.031 ± 0.006% vs. Pre-LPS: 0.041 ± 0.007%, t(20) = 1.073, p > 0.05) (Figure 9C).

**Figure 9.**
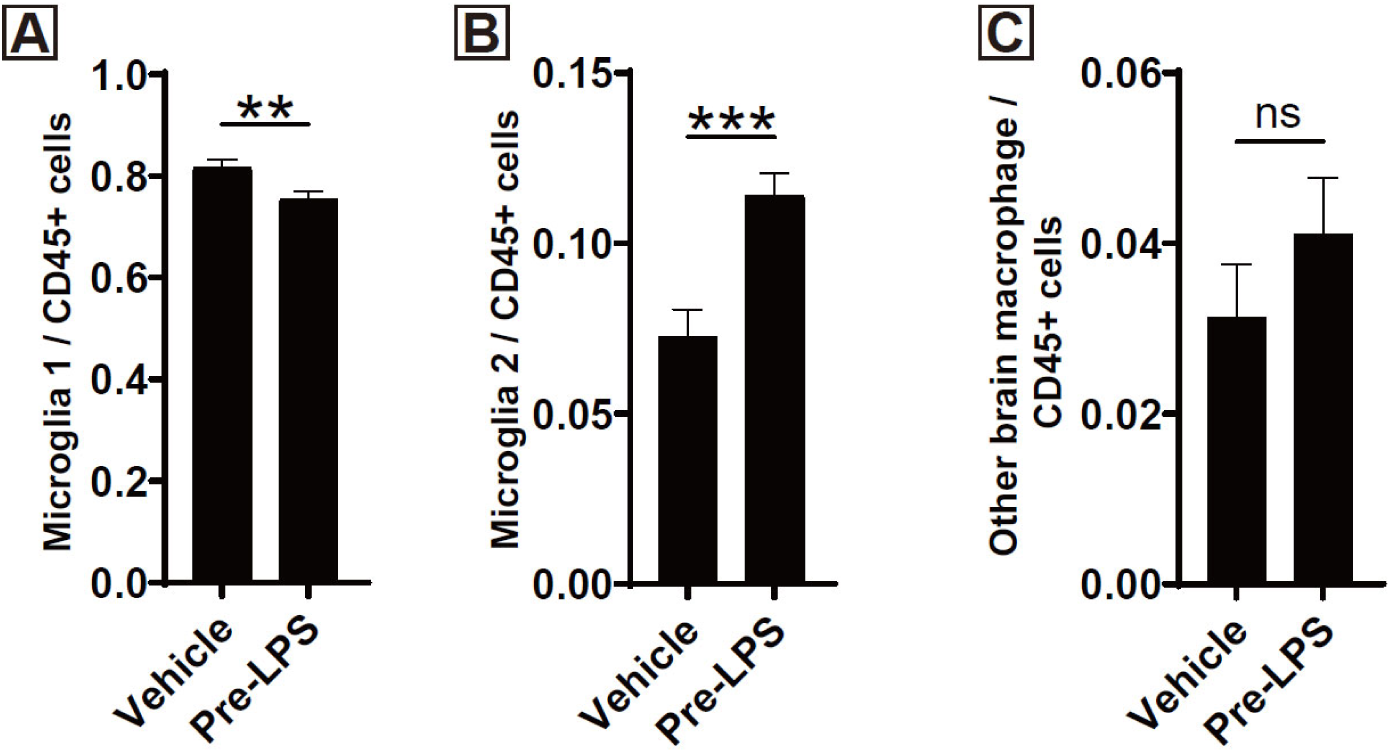
Preconditioning alters brain macrophage subpopulation proportions. (A) Microglia 1 proportion. (B) Microglia 2 proportion. (C) Other brain macrophages proportion. Unpaired t-test. n = 11 per group. **p < 0.01, ***p < 0.001. Data represent mean ± SEM.

### 5. LPS preconditioning induces subpopulation-specific transcriptional changes in brain macrophages

To examine the effects of LPS preconditioning on gene expressions in each brain macrophage subpopulation, we performed RNA-seq analysis following cell sorting by flow cytometry. Microglia 1, Microglia 2, and Other brain macrophages were isolated from Control (n=12) and Pre-LPS (n=12) groups, and transcriptional profiles were compared.

Differential gene expression analysis revealed varying degrees of transcriptional changes across subpopulations following preconditioning (Figure 10, Table 2). In Microglia 1, 2 upregulated and 5 downregulated genes were identified. Microglia 2 showed the least change with only 1 upregulated gene identified. In contrast, Other brain macrophages displayed marked transcriptional changes with 115 genes upregulated and 15 downregulated (FDR < 0.05).

**Figure 10.**
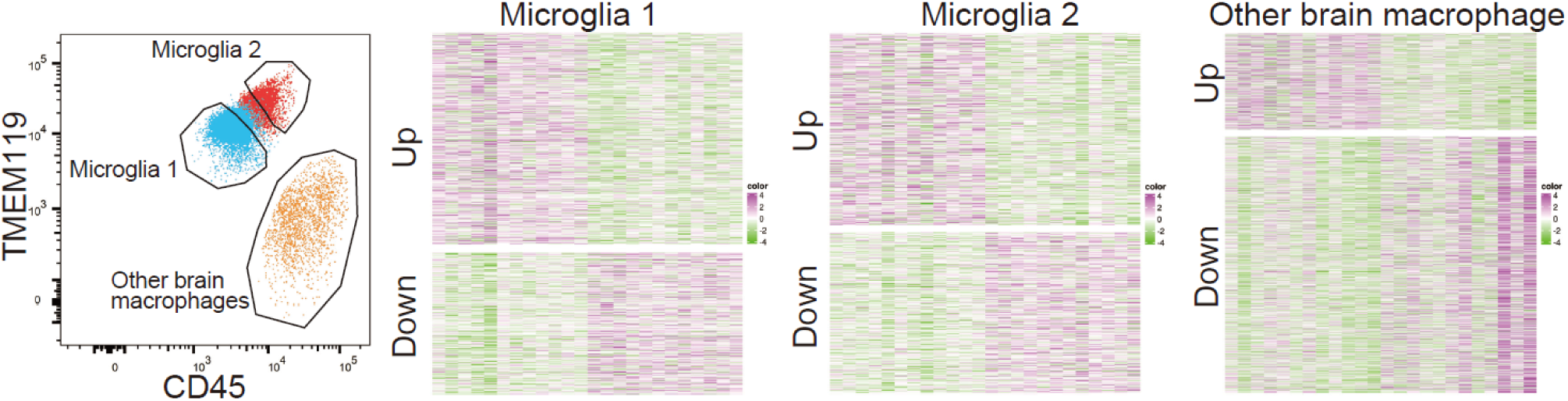
Differential gene expression analysis. Volcano plots showing differentially expressed genes in (A) Microglia 1, (B) Microglia 2, and (C) Other brain macrophages. Red dots indicate upregulated genes, blue dots indicate downregulated genes (p < 0.05, |fold change| > 1).

**Table 2.** Number of significantly upregulated and downregulated genes in each brain macrophage subtype following LPS preconditioning. Differential expression was assessed between control and LPS preconditioned groups within each cell subtype. Genes were considered significant if they met the threshold of false discovery rate (FDR)–adjusted p-value (q-value) < 0.05.

|  | Microglia 1 | Microglia 2 | Other brain macrophage |
| --- | --- | --- | --- |
| Up-regulated | 2 | 1 | 115 |
| Down-regulated | 5 | 0 | 15 |

Gene Ontology analysis revealed statistically significant functional category enrichment only in Microglia 2 (Figure 11, Table 3). Despite having the fewest differentially expressed genes, Microglia 2 showed enrichment of immune response-related pathways including response to LPS, response to molecule of bacterial origin, and regulation of DNA-binding transcription factor activity (FDR < 0.05). In Microglia 1 and Other brain macrophages, metabolic processes and inflammation-related terms were identified but did not reach statistical significance (FDR > 0.05).

**Figure 11.**
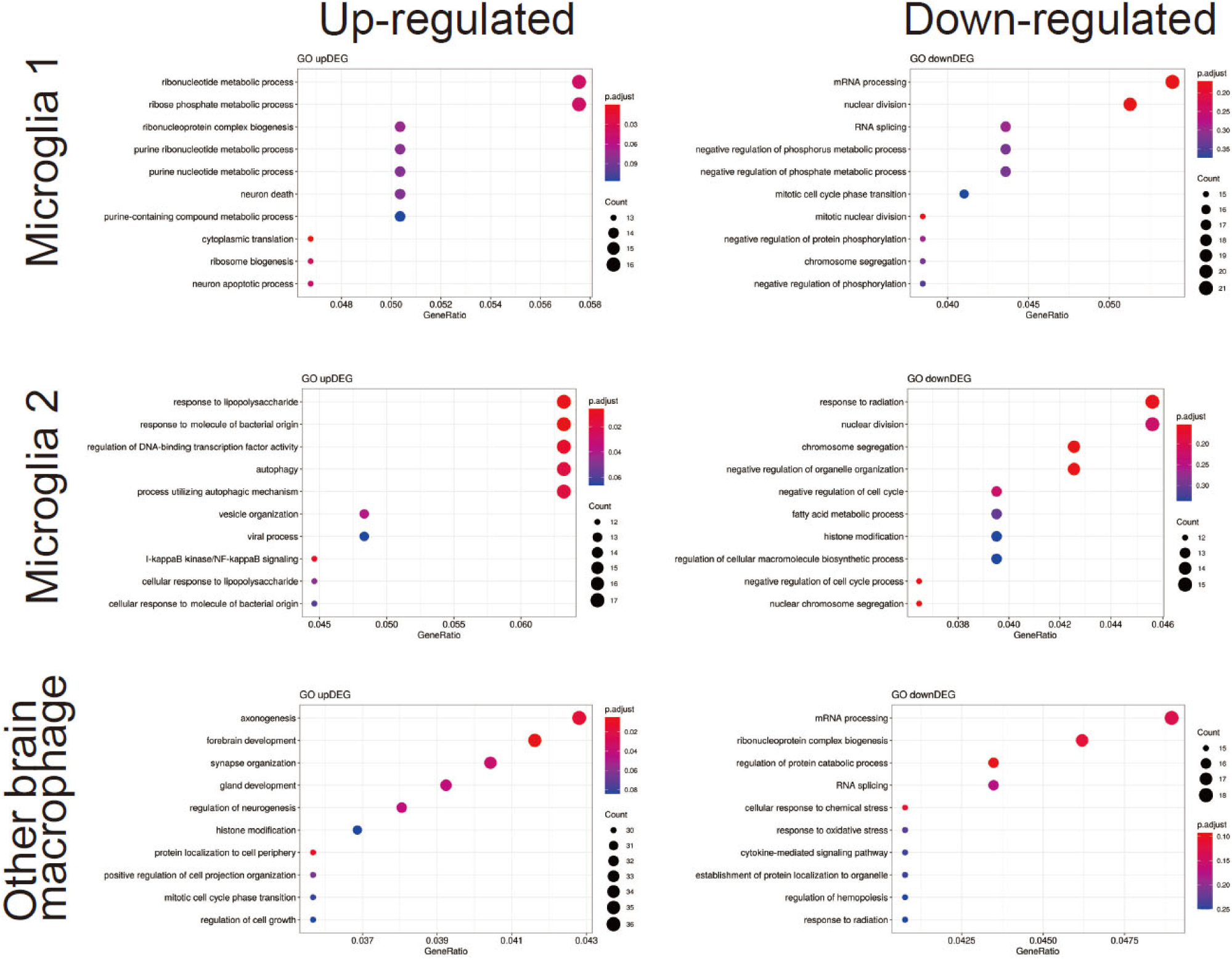
Gene Ontology enrichment analysis. Dot plots showing enriched GO terms for differentially expressed genes in each subpopulation. Dot size represents gene count, color represents adjusted p-value. Only Microglia 2 showed statistically significant enrichment (FDR < 0.05).

**Table 3.** Summary of LPS preconditioning-induced responses across brain macrophage subtypes. Qualitative transcriptional changes in three brain macrophage subtypes are summarized based on GO term enrichment and pathway activation analyses. Pathway annotations for Microglia 1 and Other brain macrophages were derived from non-significant enrichment analyses (adjusted p > 0.05) and are shown for reference only, whereas Microglia 2 exhibited statistically significant enrichment (adjusted p < 0.05).

| Cell subtype | Representative functional categories |
| --- | --- |
| Microglia 1 | Negative regulation of cell cycle<br>(*non-significant*) |
| Microglia 2 | Response to LPS, cytokine production, immune activation (*significant*) |
| Other brain macrophage | Limited oxidative stress and metabolic process terms<br>(*non-significant*) |

Pathway analysis showed similar trends, suggesting different signaling pathway changes in each subpopulation, though most did not reach statistical significance (Figure S1).

These results indicate that preconditioning primarily induced numerous gene expression changes in Other brain macrophages. While Microglia 2 had the fewest differentially expressed genes, it showed significant enrichment of immune response-related pathways.

### 6. LPS preconditioning prevents behavioral abnormalities in a chronic corticosterone model

To examine whether the protective effects of LPS preconditioning extend beyond acute systemic inflammation, we tested a chronic corticosterone administration model, which represents HPA axis hyperactivity and is widely used to induce anxiety and depressive-like behaviors. Male C57BL/6N mice received LPS (0.2 mg/kg) or saline preconditioning on Days 1-2, followed by chronic corticosterone exposure (140 μg/mL in drinking water) beginning on Day 9 for 4 weeks.

Behavioral analysis using the marble burying test revealed that corticosterone significantly increased anxiety-like behavior compared to vehicle-treated controls (Saline-Vehicle: 3.14 ± 0.99 marbles vs. Saline-COR: 9.33 ± 1.43 marbles, p < 0.01), and LPS preconditioning significantly prevented this increase (Saline-COR: 9.33 ± 1.43 marbles vs. Preconditioning-COR: 3.00 ± 1.32 marbles, p < 0.01) (Figure 12A).

**Figure 12.**
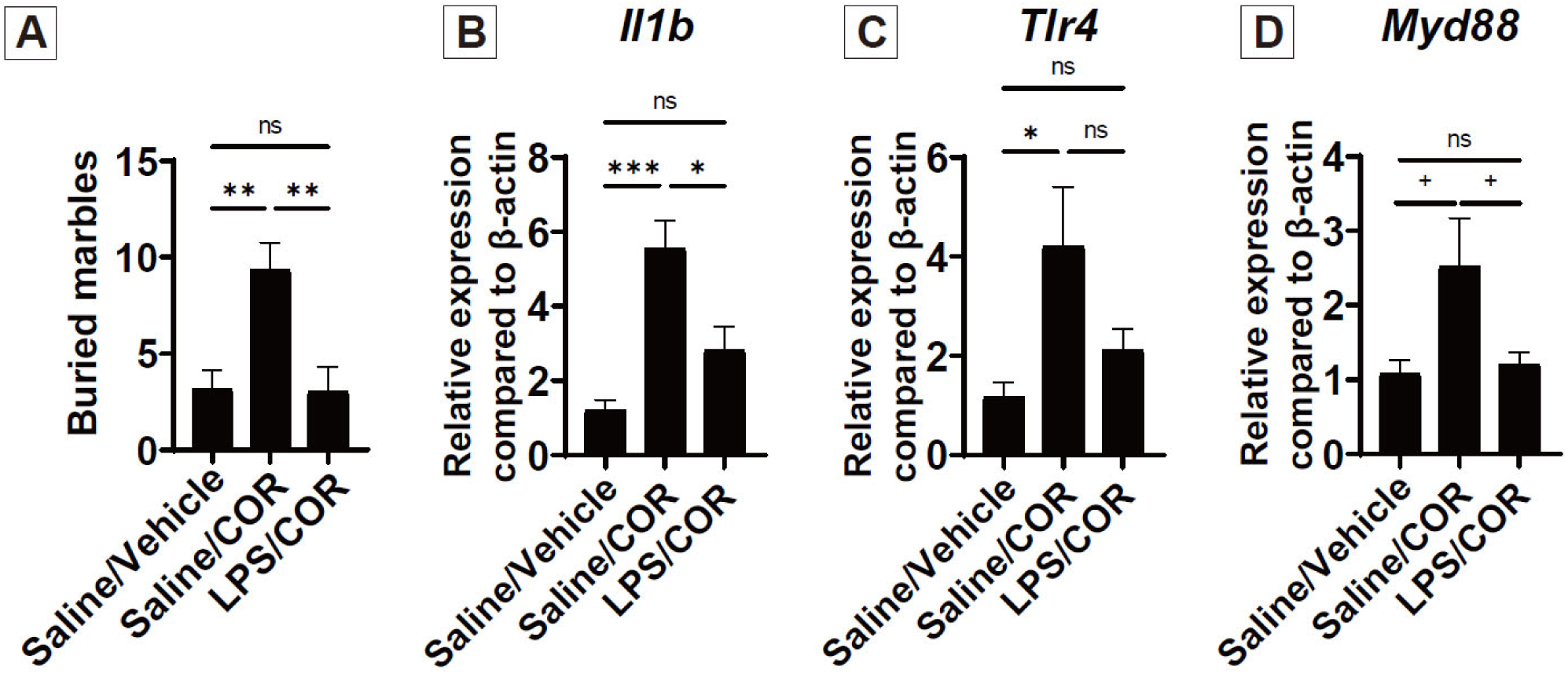
LPS preconditioning prevents behavioral abnormalities and attenuates inflammatory gene expression in a chronic corticosterone model. (A) Number of marbles buried in the marble burying test after 4 weeks of corticosterone exposure (140 μg/mL in drinking water). (B-D) Hippocampal gene expression of *Il1b*, *Tlr4*, and *Myd88* measured by qRT-PCR, normalized to *Actb* (β-actin). One-way ANOVA with Tukey’s post-hoc test. *p < 0.05, **p < 0.01, ***p < 0.001; +p < 0.1 (trend); ns, not significant. A: Saline-Vehicle (n=7), Saline-COR (n=6), LPS-COR (n=6). B: Saline-Vehicle (n=7), Saline-COR (n=6), LPS-COR (n=5). C-D: Saline-Vehicle (n=6), Saline-COR (n=6), LPS-COR (n=6). Outliers were excluded using the IQR method prior to analysis. Data represent mean ± SEM.

Hippocampal gene expression analysis demonstrated that corticosterone markedly upregulated Il1b (Saline-Vehicle: 1.2 ± 0.28 vs. Saline-COR: 5.55 ± 0.75, p < 0.001), which was significantly attenuated by preconditioning (Saline-COR: 5.55 ± 0.75 vs. Preconditioning-COR: 2.81 ± 0.63, p < 0.05). Similarly, Tlr4 expression was upregulated by corticosterone (Saline-Vehicle: 1.16 ± 0.30 vs. Saline-COR: 4.21 ± 1.20, p < 0.05), while Myd88 exhibited a trend toward suppression by preconditioning (Saline-COR: 2.52 ± 0.65 vs. Preconditioning-COR: 1.21 ± 0.16, p < 0.1) (Figure 12B-D).

These findings indicate that LPS preconditioning can prevent behavioral abnormalities and attenuate neuroinflammatory responses not only in acute systemic inflammation but also in a chronic stress model.

## Discussion

This study demonstrates that low-dose LPS preconditioning suppresses behavioral changes following systemic inflammation, particularly social behavioral deficits. This protective effect was dependent on the presence of microglia, as microglial depletion during the preconditioning period abolished the effect. Furthermore, preconditioning induced specific changes in brain macrophage subpopulation composition and transcriptional profiles.

Region-specific neuroinflammatory responses were observed between hippocampal CA1 and CA3. In CA1, systemic inflammation caused marked increases in PBR signal and PBR/IBA1 ratio, which preconditioning significantly suppressed. In contrast, CA3 showed milder changes with systemic inflammation and limited preconditioning effects. This regional specificity may reflect functional differences between regions in social behavior. CA1 is important for information integration and memory retrieval, contributing to social context recognition ^23^. In contrast, CA3 is responsible for pattern separation and novel stimulus processing ^22^. The social behavioral deficits and pronounced neuroinflammation in CA1 observed in this study suggest that social memory retrieval and integration processes may be more vulnerable to inflammation.

Immunohistochemical analysis showed that preconditioning suppressed the increase in PBR/IBA1 ratio in hippocampal CA1. PBR is known as a microglial activation marker ^33^, and these results suggest preconditioning limits excessive microglial activation. Importantly, microglial depletion experiments causally demonstrated that microglia are required for the behavioral protective effect of preconditioning.

A key feature of our experimental design was depleting microglia only during the preconditioning period, then allowing repopulation. PLX3397-mediated depletion was incomplete, with residual microglia remaining ^29^. Post-withdrawal repopulation likely occurred through division of these residual microglia ^29^. Interestingly, the loss of behavioral protective effects despite microglial repopulation indicates that the microglial state during preconditioning induction is crucial for subsequent effect manifestation.

Epigenetic regulation has been implicated in endotoxin tolerance maintenance mechanisms. DNA methylation-mediated suppression of inflammation-related genes^34, 35^ and chromatin remodeling-mediated transcriptional control^36^are known. Given that DNA methylation states are maintained by DNMT1 after cell division^37, 38^, while chromatin modifications are not necessarily stably inherited ^39, 40^, DNA methylation-mediated regulatory mechanisms may be involved in this study.

Flow cytometry and RNA-seq analyses revealed brain macrophage heterogeneity and preconditioning-specific responses. Using our previously reported TMEM119/CD45 expression pattern classification^28^, we observed decreased proportions of TMEM119-positive CD45-low cells (defined as Microglia 1 in this study) and increased TMEM119-positive CD45-intermediate cells (defined as Microglia 2). These changes suggest that preconditioning alters the balance between microglial subpopulations, with each acquiring different functional states.

Interestingly, transcriptional change patterns differed markedly among subpopulations following preconditioning. Despite having only one differentially expressed gene, Microglia 2 showed significant enrichment of LPS response-related pathways, suggesting functionally important though limited changes. Meanwhile, Microglia 1 had seven differentially expressed genes identified but no significant functional category enrichment.

The lack of specific functional category enrichment despite extensive transcriptional changes (130 differentially expressed genes) in TMEM119-negative CD45-high cells (defined as Other brain macrophages) may reflect this population’s heterogeneity. This population includes border-associated macrophages (BAMs) localized to different anatomical sites such as perivascular, meningeal, and choroid plexus macrophages ^41, 42^. Individual transcriptional responses of these to preconditioning may have resulted in no unified functional changes detected in the overall population. Future detailed analysis using single-cell RNA-seq is needed to clarify individual roles of these subpopulations.

Importantly, we demonstrated that the protective effects of microglial preconditioning extend beyond acute inflammatory challenges to include chronic stress-induced behavioral abnormalities. In the chronic corticosterone administration model, LPS preconditioning prevented anxiety-like behavior and attenuated hippocampal inflammatory gene expression. The suppression of Il1b upregulation is particularly noteworthy, as this pro-inflammatory cytokine has been implicated in anxiety and depression-like behaviors (PMID: 22386198; PMID: 26711676). These findings suggest a broad spectrum of protection conferred by microglial preconditioning against diverse neuropsychiatric symptom-inducing stimuli with inflammatory components.

## Study Limitations

Several limitations should be acknowledged. First, LPS administration at the doses used in this study resulted in some animal mortality, which is an inherent limitation of this model. While this reduced our final sample sizes, the remaining animals still provided sufficient statistical power to detect the primary outcomes.

Second, this study used only male mice. While sex differences in immune responses and behavioral changes are an important research topic ^43^, we limited to males to exclude confounding factors such as estrous cycle in this initial investigation. Future studies in females are needed.

Third, how microglial changes modulate neural circuit activity leading to behavioral changes remains unclear. Effects on synaptic function and neuronal activity require investigation.

Fourth, detailed identification of epigenetic regulatory mechanisms is needed. Whether DNA methylation or chromatin remodeling plays the primary role, and identification of target genes, remain future challenges.

## Conclusion

This study is the first to demonstrate the possibility of preventive intervention for neuroinflammation-induced behavioral abnormalities. It suggests that immune regulation targeting microglia could be a preventive strategy for psychiatric symptoms, providing a new perspective on the neuroinflammation hypothesis. Particularly, the discovery of microglial subpopulation-specific responses emphasizes the importance of targeted interventions for specific cell populations rather than uniform anti-inflammatory treatment. The failure to maintain preconditioning effects after microglial turnover indicates the importance of cellular state during preconditioning induction while suggesting the existence of immune memory mechanisms via epigenetic regulation. Future detailed elucidation of molecular mechanisms is expected to lead to development of novel preventive and therapeutic strategies for psychiatric disorders.

## Data availability

RNA-sequencing data from 72 samples (3 brain macrophage subpopulations × 2 conditions × 12 biological replicates) have been deposited in the Gene Expression Omnibus (GEO). Deposited data include raw FASTQ files and normalized expression matrices (CPM) for each subpopulation comparison. All other data supporting the findings of this study are available from the corresponding author upon reasonable request.

## Code availability

RNA-seq analysis was performed using the standard pipeline implemented by cBioinformatics Inc., utilizing edgeR (v3.40.2), gprofiler2 (v0.2.1), clusterProfiler (v4.6.2), and RcisTarget (v1.18.2) as described in Methods. No custom code was generated for this study.

## Author contributions

M.K (Minori). conceived the study, designed and performed experiments, analyzed data, and wrote the manuscript. H.N., M.S. (Masafumi), and M.K. (Manabu) performed flow cytometry and cell sorting experiments. M.S. (Mayumi), R.N., T.S., and F.A. performed behavioral experiments and immunohistochemistry. T.I. provided technical assistance. M.K. (Manabu) contributed to study design. H.T. supervised the study and reviewed the manuscript. All authors approved the final manuscript.

## Declaration of generative AI in scientific writing

The authors used Claude (Anthropic) to improve the clarity and grammatical accuracy of the English language in this manuscript and to generate illustrations for Figure 1 using ChatGPT (DALL·E), OpenAI. All scientific content, experimental design, data analysis, and interpretation are the original work of the authors.

## Acknowledgments

We thank the staff of the Laboratory Animal Resource Center at National Defense Medical College for animal care.

