## Supplementary materials for "Microglia-dependent LPS preconditioning prevents neuroinflammation-induced behavioral deficits in male mice"

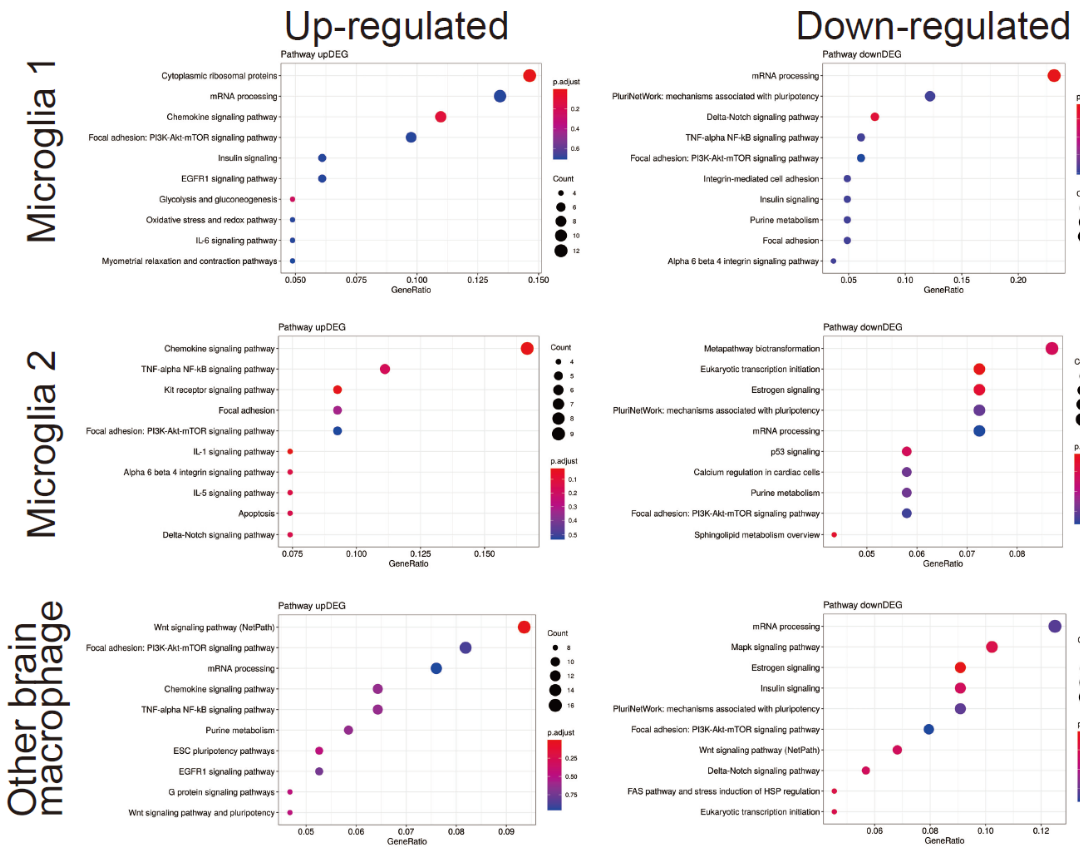


**Supplementary figure 1**

Pathway enrichment analysis across brain macrophage subpopulations

Dot plots showing pathway enrichment for upregulated (upper panels) and downregulated genes (lower panels) in Microglia 1, Microglia 2, and Other brain macrophages following LPS preconditioning. Dot size represents gene count, color represents adjusted p-value. Most pathways did not reach statistical significance (adjusted $p > 0.05$).

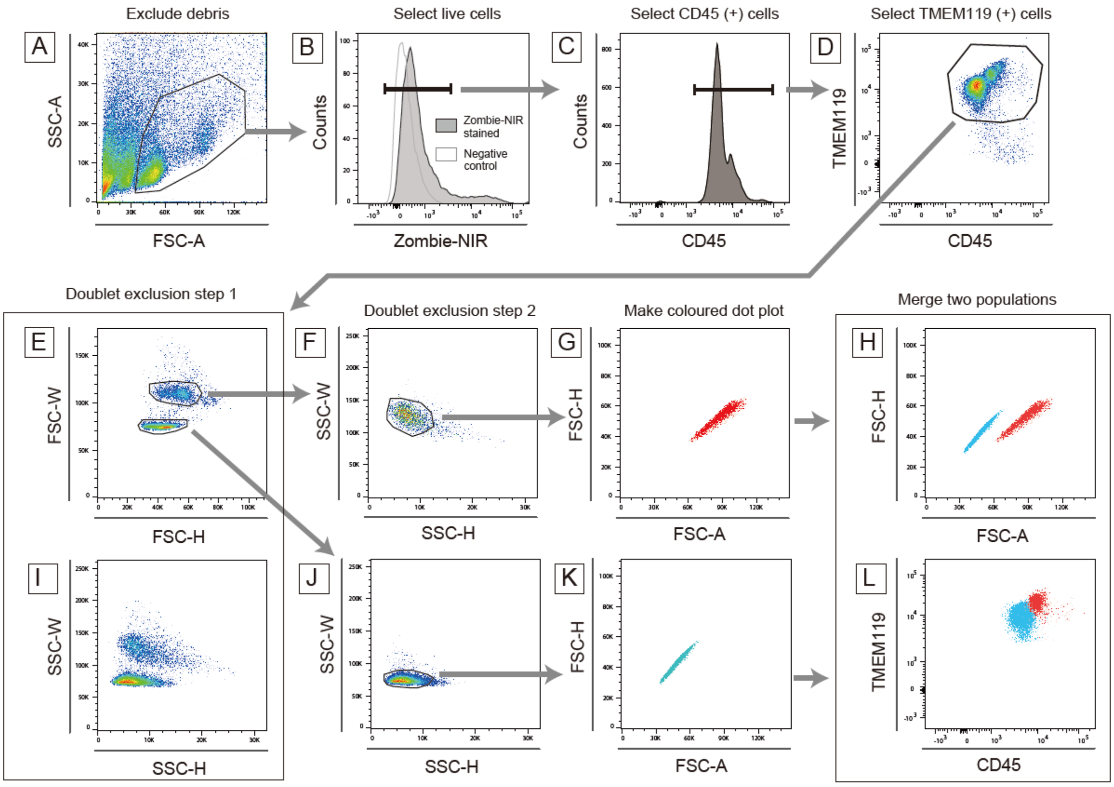


**Supplementary figure 2**

Live-dead analysis and doublet exclusion of Microglia 1 and 2.

To ensure that Microglia 1 and 2 are distinct populations, we employed live-dead marker Zombie NIR staining and carefully excluded doublet cells using a two-step gating strategy based on forward scatter (FSC) and side scatter (SSC) analysis. (A) The extracted CD11b-positive cells were subjected to flow cytometry. Debris was excluded, and an appropriate cell size population was gated using the FSC area (FSC-A) / SSC area (SSC-A) histogram. (B) The dead cell marker Zombie NIR was deployed to distinguish between live and dead cells. Comparing negative control staining, Zombie negative cells were identified as live cells and selected for analysis. (C) Most gated cells were positive for CD45 and were gated for further study. (D) TMEM119 and CD45 double-positive cells were microglial cells, and (E) the gated TMEM119 positive cells were developed into a forward scatter height (FSC-H) and width (FSC-W) histogram. Two distinct populations were observed, with the signal strength of FSC-W being significantly different and without overlap. This histogram revealed two distinct populations with different cell sizes. A major part of each population was selected as single cells (Step 1). (I) The SSC height (SSC-H) and width (SSC-W) histograms of the same population demonstrated that the two distinct populations had different cellular complexities, which was consistent with the data from the FSC-H and FSC-W histograms. (F, J) Each selected single cell (Step 1) was developed into SSC-H and SSC-W, and significant populations were gated as single cells (Step 2). (G, K) Each single cell population was represented as a dot plot histogram, with FSC-A and FSC-H displayed in different colors. (H) The two histograms were merged into a single one, which showed that two distinct populations exist based on their cellular structures. (L) The CD45 and TMEM119 histogram showed that these two populations have different expression levels of each antigen.
